# A modeling analysis of whole-body potassium regulation on a high potassium diet: Proximal tubule and tubuloglomerular feedback effects

**DOI:** 10.1101/2023.12.12.571254

**Authors:** Melissa M. Stadt, Anita T. Layton

## Abstract

Potassium (K^+^) is an essential electrolyte that plays a key role in many physiological processes, including mineralcorticoid action, systemic blood-pressure regulation, as well as hormone secretion and action. Indeed, maintaining K^+^ balance is critical for normal cell function, as too high or too low K^+^ levels can have serious and potentially deadly health consequences. K^+^ homeostasis is achieved by an intricate balance between the intracellular and extracellular fluid as well as balance between K^+^ intake and excretion. This is achieved via the coordinated actions of regulatory mechanisms such as the gastrointestinal feedforward effect, insulin and aldosterone upregulation of Na^+^-K^+^-ATPase uptake, and hormone and electrolyte impacts on renal K^+^ handling. We recently developed a mathematical model of whole-body K^+^ regulation to unravel the individual impacts of regulatory mechanisms. In this study, we extend our mathematical model to incorporate recent experimental findings that showed decreased fractional proximal tubule reabsorption under a high K^+^ diet. We conducted model simulations and sensitivity analyses to unravel how these renal alterations impact whole-body K^+^ regulation. Our results suggest that the reduced proximal tubule K^+^ reabsorption under a high K^+^ diet could achieve K^+^ balance in isolation, but the resulting tubuloglomerular feedback reduces filtration rate and thus K^+^ excretion. Model predictions quantify the sensitivity of K^+^ regulation to various levels of proximal tubule K^+^ reabsorption adaptation and tubuloglomerular feedback. Additionally, we predict that without the hypothesized muscle-kidney cross talk signal, intracellular K^+^ stores can exceed normal range under a high K^+^ diet.

## 1 Introduction

Potassium (K^+^) homeostasis is critical for normal cell function. The function of excitable cells is controlled by membrane voltage, which is primarily determined by the ratio of the extra- and intracellular K^+^ concentrations ([K^+^]) [44]. K^+^ also plays a key role in other physiological processes such as mineralcorticoid action, systemic blood-pressure regulation, as well as hormone secretion and action [14, 23, 25, 29].

About 98% of total body K^+^ is stored in the intracellular fluid, primarily within the skeletal muscle cells [8, 23]. This results in a concentration gradient between the intracellular and extracellular fluid with a high intracellular [K^+^] (120-140 mmol/L) and a low extracellular [K^+^] (3.5-5.0 mmol/L) [8, 14, 23]. In total this means the amount of K^+^ in the extracellular fluid is around 65 mmol, which can easily be exceeded by dietary K^+^ intake. This homeostatic challenge is overcome by regulatory mechanisms such as insulin signalling and Na^+^-K^+^-ATPase uptake that moves excess extracellular K^+^ into the intracellular fluid. In a healthy person, the kidneys ensure that K^+^ excretion is balanced with K^+^ intake [5, 15, 18]. K^+^ homeostasis is achieved by balancing the extra- and intracellular distribution as well as balancing excretion with intake, primarily via urinary excretion. When one of these mechanisms are altered, hyper- or hypokalemia may occur. Due to the importance of K^+^, and the dangers of hyper- and hypokalemia, humans have evolved to have several regulatory mechanisms that ensure extracellular [K^+^] stays within a normal range.

For much of the evolution of the K^+^ regulatory systems, humans consumed a diet rich in fruits, vegetables, nuts, and seeds. This diet contained ample amounts of K^+^ and very little Na^+^. Specifically, it is estimated that in Paleolithic times, the average intake for K^+^ was about 230-400 mmol/day, which is about four-fold the amount of K^+^ in the modern Western diet (about 70-100 mmol/day) due to significant changes in food sources [8, 14]. This change from a K^+^-rich diet to a K^+^-poor diet (in Western cultures) is estimated to have occurred within the last 5,000-10,000 years, which is a short time frame from an evolution perspective. Therefore, it is likely that our K^+^ regulatory systems are better suited for a K^+^-rich diet [4, 14, 29]. This may explain why the modern high Na^+^, low K^+^ diet is associated with many health problems such as high blood pressure, cardiovascular disease, and strokes [4, 14, 25, 29]. Notably, it has been found that the Yanomamö natives in the Amazon, who are known to have a K^+^-rich, low Na^+^ diet, closer to that of Paleolithic times, do not have the same increases in blood pressure as they age in contrast to Western cultures [27].

To unravel the complexities of K^+^ regulation, we developed a detailed compartmental model of whole-body K^+^ regulation in a recent study (Ref. [39]). The model, which consists of a system of coupled ordinary differential equations (ODE) and algebraic equations, represents the intra- and extracellular fluid compartments as well as a detailed renal compartment. The model simulates known regulatory mechanisms: (i) the gastrointestinal feedforward control mechanism, (ii) the effect of insulin and (iii) aldosterone (ALD) on cellular K^+^ uptake via Na^+^-K^+^-ATPase, and (iv) ALD stimulation of renal K^+^ secretion. We used this model to quantify the impacts of individual feedback and feedforward effects. In our previous study, we found that the model predicted high extra- and intracellular [K^+^] under high K^+^ intake despite it being known that healthy persons can sustain a high K^+^ diet [32]. It is likely that there are other mechanisms of K^+^ handling during a high K^+^ diet that have not been fully understood. In our previous study, we proposed an additional hypothesized regulatory mechanism, muscle-kidney cross talk, where [K^+^] in the skeletal muscle cells directly affect renal K^+^ transport without changes in extracellular [K^+^] [39]. This signal would enable mammals to sustain normal levels of plasma and intracellular [K^+^] during high K^+^ intake.

A recent experimental study in rats showed evidence for inhibited proximal tubule (PT) K^+^ reabsorption under a high K^+^ diet when compared to controls (Ref. [42]). The authors also showed decreased glomerular filtration rate (GFR) due to the tubuloglomerular feedback (TGF) signal from inhibited Na^+^ and K^+^ reabsorption along the PT in rats on a high K^+^ diet [42].

In this study, we extend our mathematical model to include the impact of a high K^+^ diet on PT K^+^ reabsorption as well as TGF effects. The objectives of this study are to: (i) update our mathematical model of whole-body K^+^ regulation to include new experimental findings, (ii) quantify how renal adaptations of PT K^+^ reabsorption and TGF effects impact whole-body potassium homeostasis, (iii) investigate whole-body impacts of varied renal adaptations under a high K^+^ diet, and (iv) conduct *in silico* experiments of the muscle-kidney cross talk signal on our updated model.

## 2 Methods

In this study, we investigate the PT and TGF effects of high K^+^ intake reported by Wang et al. [42]. We use our previously developed mathematical model of whole-body K^+^ regulation (Ref. [39]) with modifications described below. A model schematic is given in Fig. 2.1 and a more detailed model description is available in Appendix B.

**Figure 2.1:**
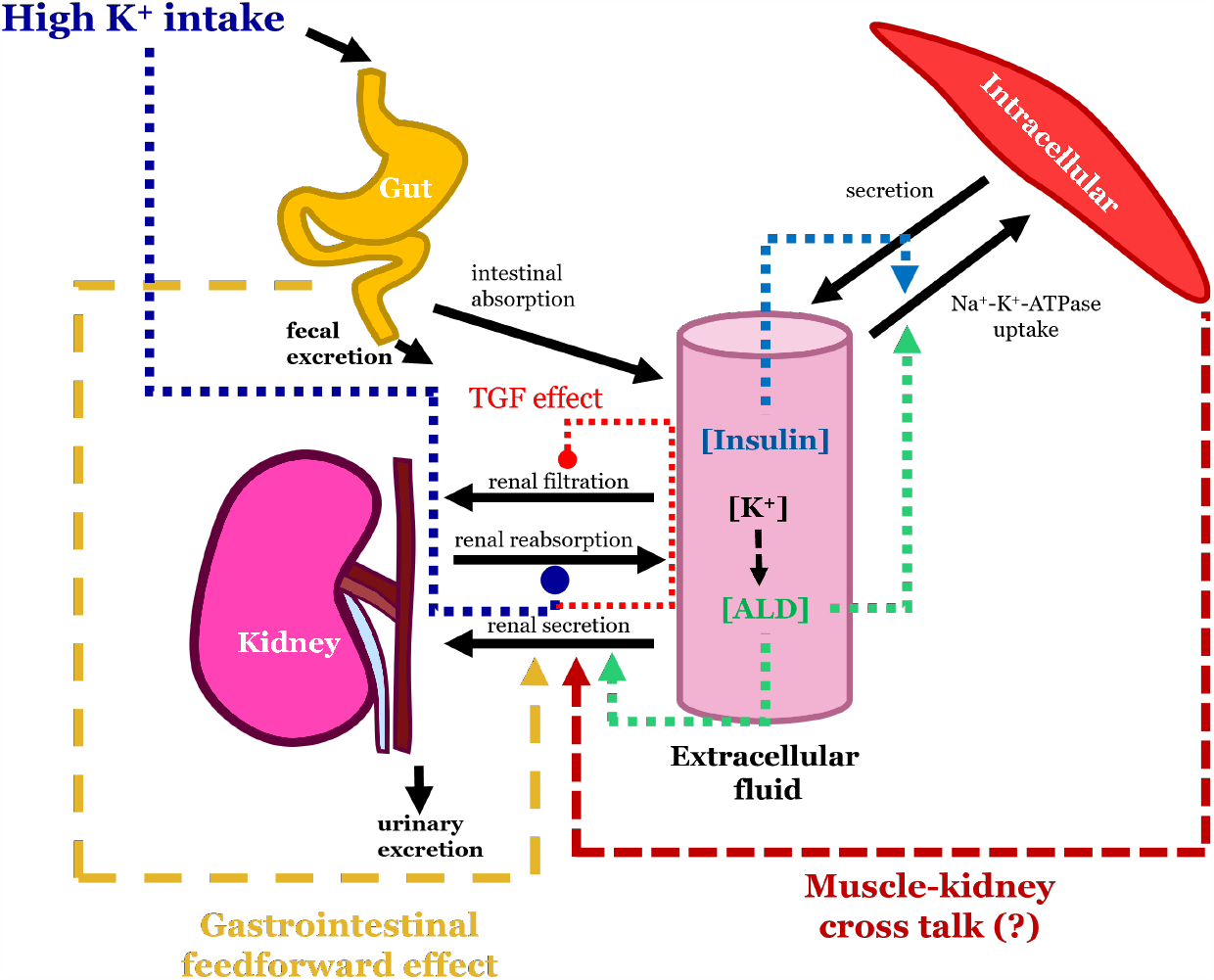
Model schematic under high K^+^ intake. Black arrows represent transport of K^+^ between the compartments. Green, light blue, and yellow arrows represent the stimulating effects of aldosterone, insulin, and the gastrointestinal feedforward mechanism, respectively. High K^+^ intake inhibitory effects on the proximal tubule (i.e., renal reabsorption) reabsorption is shown by the blunted dark blue arrow. Tubuloglomerular feedback (TGF) is shown by the blunted light red arrow. The hypothesized muscle-kidney cross talk signal is shown by the dark red arrow. The extracellular compartment is made up of plasma and interstitial fluid. ALD: aldosterone; TGF: tubuloglomerular feedback

**Figure 2.2:**
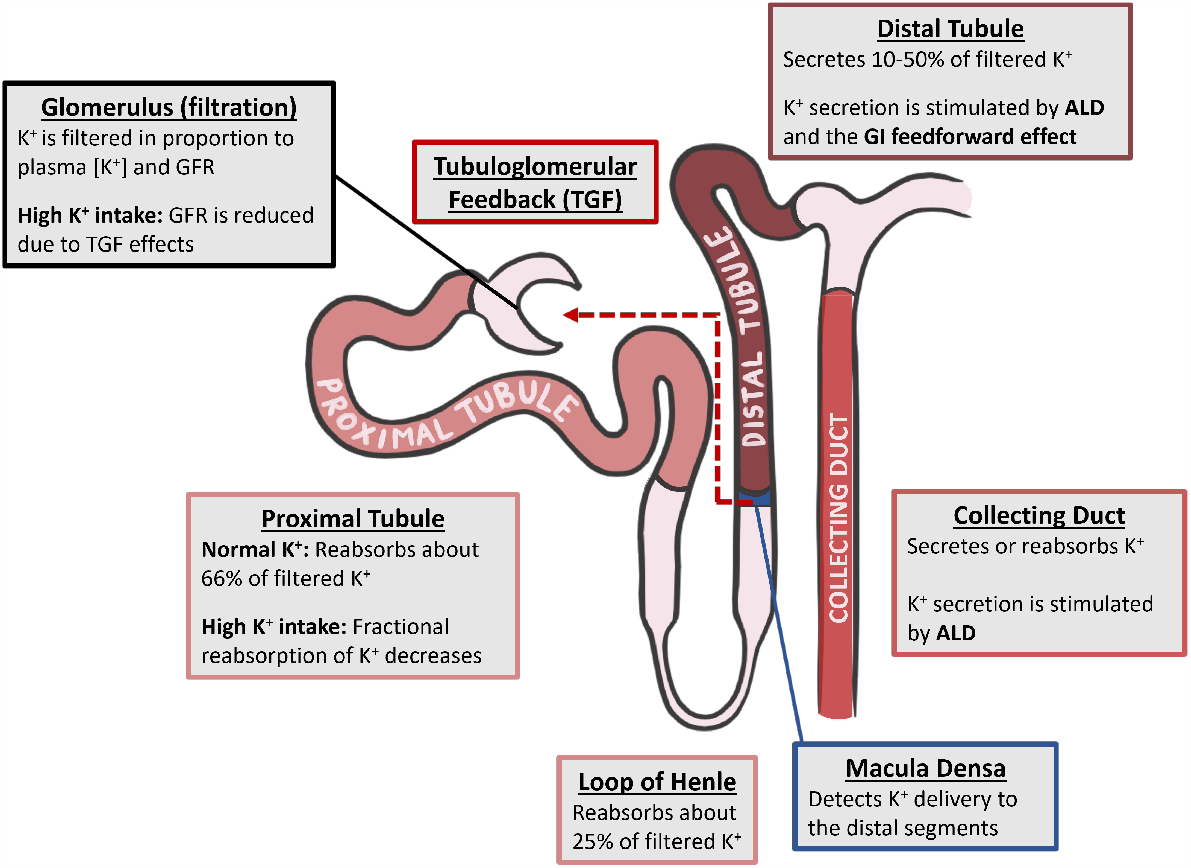
Schematic of renal K^+^ handling along various nephron segments in the kidneys. ALD: aldosterone; GFR: glomerular filtration rate; TGF: tubuloglomerular feedback

### 2.1 High K^+^ diet simulation

To conduct simulations of a control or chronic high K^+^ diet in a human (man), we added a fixed amount of K^+^ to the gut compartment at meal times for 50 days. We simulated 3 meals of equal content per day at the simulation hour 6, 12, and 18. For the control K^+^ diet, the 3 meals total 78 mmol per day, typical of an average modern Western diet [23]. For the high K^+^ diet, we increased the K^+^ intake per meal by four folds, totalling 312 mmol per day. This diet protocol is proportional to the control and high K^+^ diet experimental protocols conducted in rats in Ref. [42]; more below.

### 2.2 Impact of high K^+^ intake on proximal segment K^+^ reabsorption

In a recent experimental study, Wang et al. [42] showed that rats fed a high K^+^ diet for 7 days exhibited decreased PT fractional K^+^ reabsorption and subsequent reduction in GFR. This is associated with a significant increase in urinary K^+^ excretion to balance the high K^+^ load. Specifically, Wang et al. [42] reported that fractional PT K^+^ reabsorption in rats on a high K^+^ diet was 37% versus 66% in control diet rats. Similar changes occurred for PT fractional Na^+^ reabsorption [42]. Additionally, the authors reported the GFR decreased by 29% as a result of TGF activated by the decreased delivery of NaCl and K^+^ to the macula densa.

To investigate the impacts of a high K^+^ diet on whole-body K^+^ regulation, we incorporate into our model the effects of a high K^+^ diet on fractional PT K^+^ reabsorption and TGF. Let *η*_pt−Kreab_ denote the fractional K^+^ reabsorption along the PT. Under normal K^+^ intake this is about 67% [5, 39, 42]. Fractional reabsorption of K^+^ along the loop of Henle is denoted by *η*_LoH−Kreab_ = 25%. Let “proximal segment” refer to the segments before the macula densa (i.e., the PT and loop of Henle) so that

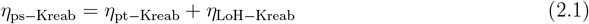

*η*_ps−Kreab_ = *η*_pt−Kreab_ + *η*_LoH−Kreab_ (2.1) where *η*_ps−Kreab_ denotes the total proximal segment fractional reabsorption.

To simulate the TGF effect of altered fractional PT K^+^ reabsorption, we determine GFR (Φ_GFR_) by

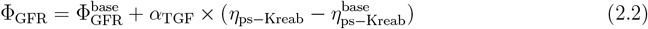

where 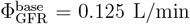 is the baseline GFR, 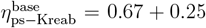 is the baseline fractional proximal segment reabsorption, and *α*_TGF_ is a parameter representing the strength of the TGF signal. We refer to this equation as the “TGF effect”. We compute the baseline value for *α*_TGF_ by letting 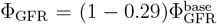 and *η*_pt−Kreab_ = 0.37 as found for the GFR and fractional PT K^+^ reabsorption in Ref. [42] under a high K^+^ diet. Therefore, plugging these values in and solving Eq. 2.2 we get that

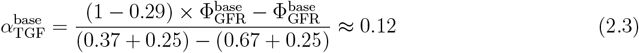

where 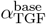 is the baseline value for *α*_TGF_. Unless otherwise noted, this is the parameter value used for simulations involving TGF effects (Eq. 2.2).

Since K^+^ is freely filtered in the glomerulus, filtered K^+^ load (Φ_filK_) is proportional to Φ_GFR_ and plasma [K^+^] (*K*_plasma_) so that

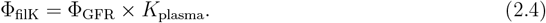

Net proximal segment K^+^ reabsorption (Φ_ps−Kreab_) is then given by

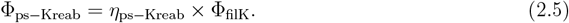

After the proximal segment, the filtrate enters the distal segments. No changes were made in this study to the mathematical modeling of distal segment K^+^ handling. A model schematic for the renal compartment is shown in Fig. 2.2. We direct readers to Ref. [39] or Appendix B for details on the model representation of these segments.

### 2.3 Sensitivity analysis

Like many models of complex physiological systems, the present whole-body K^+^ regulation model involves a large number of parameters, some of which are not measurable directly. Sensitivity analysis is a computational method that allows us to evaluate the sources of uncertainty, and to unravel the relationship between the model parameters and results [31, 33, 35]. This method has been applied to many applications of biological systems including blood clotting [20], cancer modeling [2, 33], biochemical models [13], epidemiology [21, 46], and more.

There are two primary types of sensitivity analysis: local and global methods. Local methods examine sensitivity of model behaviors to one parameter at a specific point in the parameter space. This is done by changing one parameter, fixing all other parameters at the baseline value, and evaluating the change in the model output. While this method is easy to implement and, relatively speaking, not computationally expensive, it is limited in that it does not consider interactions with other parameter changes or different points in the parameter space. In contrast, global methods compute sensitivities at multiple points and involve changing multiple parameters, thus capturing the model sensitivity to individual parameter changes, interactions with other parameters, and evaluates the model at many points in the parameter space. Global analysis can give a more comprehensive understanding of how each of the parameters in the system impact the model output. The challenge of global methods is they are typically computationally expensive, especially for models with lots of parameters. For more details about sensitivity analysis (both local and global methods), see Refs. [31, 33, 35].

Given the many parameters in the present model, a global analysis involving all the parameters is not computational feasible. As such, we seek to reduce computation time by first conducting a local sensitivity analysis to identify parameters that the model is more sensitive to, and then only include those parameters in the global sensitivity analysis. Specifically, in the first step, we conduct a local sensitivity analysis by increasing and decreasing individual parameters by 10%, and then computing the resulting plasma and intracellular [K^+^] at the end of the 50 day simulation with that individual parameter change. All other parameters remain at their baseline value. The local sensitivity is determined by

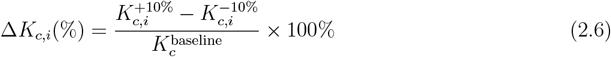

where *c* denotes the compartment (i.e., *c* = plasma or intracellular) and *i* is the parameter number. 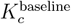 denotes the baseline [K^+^] in compartment *c* at the end of the 50 day simulation for no changes in any of the parameter values. 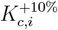 denotes the [K^+^] in compartment *c* at the end of the 50 day simulation for parameter *i* increased by 10% from its baseline value, and conversely 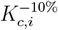 is the [K^+^] at the end of the 50 day simulation for parameter *i* decreased by 10%. Parameters that result in Δ*K*_*c,i*_ greater than 2% are included in the global sensitivity analysis.

We use the Morris method to evaluate the global sensitivity of the whole-body K^+^ regulation model under high K^+^ intake. The Morris method involves computing elementary effects for each of the parameters, which quantify how much a change in that parameter impacts the model output. Each parameter has *r* elementary effects computed at different points in the parameter space. The Morris method is known to be much more computationally efficient than other methods, allowing for parameter ranking of many parameters [31]. A more detailed description is given in Appendix C.1.

After computing the *r* elementary effects for each parameter, denoted as parameter *i*, we compute the average value (*μ*_*i*_), average of the absolute values 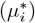, the standard deviation (*σ*_*i*_), and Morris Index (*MI*_*i*_) in the following ways:

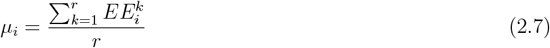

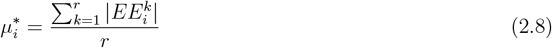

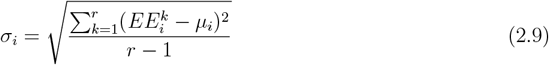

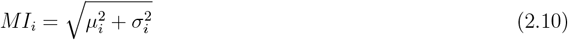

where 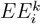 denotes the elementary effect of the *i*-th parameter during the *k*-th model evaluation. The greater the value of 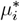 for parameter *i*, the more the *i*-th parameter affects the model output. The greater the *σ*_*i*_ value, the more the parameter is nonlinear or involved with interactions with other parameters; a low *σ*_*i*_, by contrast, indicates a linear, additive factor. We will refer to our sensitivity analysis using the Morris method as a “Morris analysis” in short.

We conduct the Morris analysis for the whole-body K^+^ regulation by sampling over the parameter space, integrating the system for a 50 day simulation under high K^+^ intake with the PT and TGF effects, and computing the final plasma and intracellular [K^+^]. The model output used for the computation of the elementary effects is the final plasma and intracellular [K^+^]. We compute *r* = 100 elementary effects for each parameter that is considered in the Morris analysis. The ranges for the parameters used are listed in Appendix C.

### 2.4 Muscle-kidney cross talk effects

In our previous study [39], we explored a hypothesized signal called “muscle-kidney cross talk” and predicted its effects under high and low K^+^ intake. This signal is between intracellular fluid (driven by skeletal muscle) and kidneys, where the skeletal muscle intracellular [K^+^] directly affects urine K^+^ excretion without changes in the extracellular [K^+^]. In the same way as Ref. [39], muscle-kidney cross talk is not included in the baseline model because there has not been definitive evidence for this signal.

In this study we further investigate the impacts of muscle-kidney cross talk in the context of a high K^+^ diet by conducting “what-if” simulations, in addition to the baseline model simulations. The coupling function (*ω*_Kic_) is taken to be a linearly increasing function of *K*_IC_, bounded above 0:

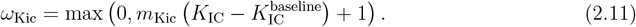

where the baseline value for *m*_Kic_ = 0.1. At homeostasis, 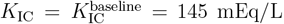, thus *ω*_Kic_ = 1. To model a muscle-kidney cross talk signal that targets distal tubule K^+^ secretion, we add the impact of *ω*_Kic_ by replacing the equation for distal segment K^+^ secretion with

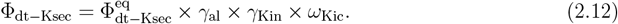

(We have previously found that the distal tubule target resulted in similar results to collecting duct reabsorption [39].)

### 2.5 Software

The simulations were conducted using `MATLAB R2022a`. Numerical solutions of the ODE system were solved using `ode15s`, a variable-step, variable-order ODE solver. Steady state solutions were obtained using the root finding function `fsolve`. For the Morris analysis we used the packages `sensitivity` and `desolve` in `R` [11, 12]. All code for the model simulations and sensitivity analysis can be found at https://github.com/Layton-Lab/highK.

## 3 Results

### 3.1 High K^+^ intake results in high intracellular K^+^

How does a high K^+^ diet impact whole-body K^+^ regulation? To investigate this question, we conducted simulations of 50 days of a normal and a high K^+^ diet (see Section 2.1) with and without different PT and TGF effects (see Section 2.2). Specifically, the high K^+^ diet simulations were conducted with:

i. no PT and TGF effects, i.e., *η*_pt−Kreab_ = 0.67 (baseline value) and *α*_TGF_ = 0. This is labeled as “High K^+^ - no PT/TGF effect” in the corresponding figures.
ii. with the PT and TGF effect found in Ref. [42], i.e., *η*_pt−Kreab_ = 0.37 and 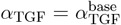. This is labeled as “High K^+^ - PT + TGF effects” in the corresponding figures.
iii. only the decrease in PT fractional reabsorption, i.e., *η*_pt−Kreab_ = 0.37 and *α*_TGF_ = 0 in Eq. 2.2. This is labeled as “High K^+^ - only PT effect” in the corresponding figures.

All simulations in this section were conducted without the muscle-kidney cross talk signal described in Section 2.4.

For the control K^+^ diet simulations, total daily K^+^ intake is 78 mmol. The model assumes that 10% of the K^+^ intake is excreted through feces [37]. Daily cumulative urinary K^+^ excretion is predicted to be 70.2 mmol (Fig. 3.1A), which is equal to 90% of the total daily K^+^ intake. Therefore, the control K^+^ diet results in balance between daily K^+^ intake and excretion. The predicted urinary K^+^ excretion corresponds to a mean fractional K^+^ excretion of 11% (Fig. 3.1B), which is within the range reported by Wang et al. [42]. Thus, on a control diet, our mathematical model captures renal handling that represents K^+^ balance as is found physiologically. This balance between daily intake and excretion results in stable total body K^+^ so that plasma and intracellular [K^+^] levels remain stable throughout the full simulation (Fig. 3.2).

**Figure 3.1:**
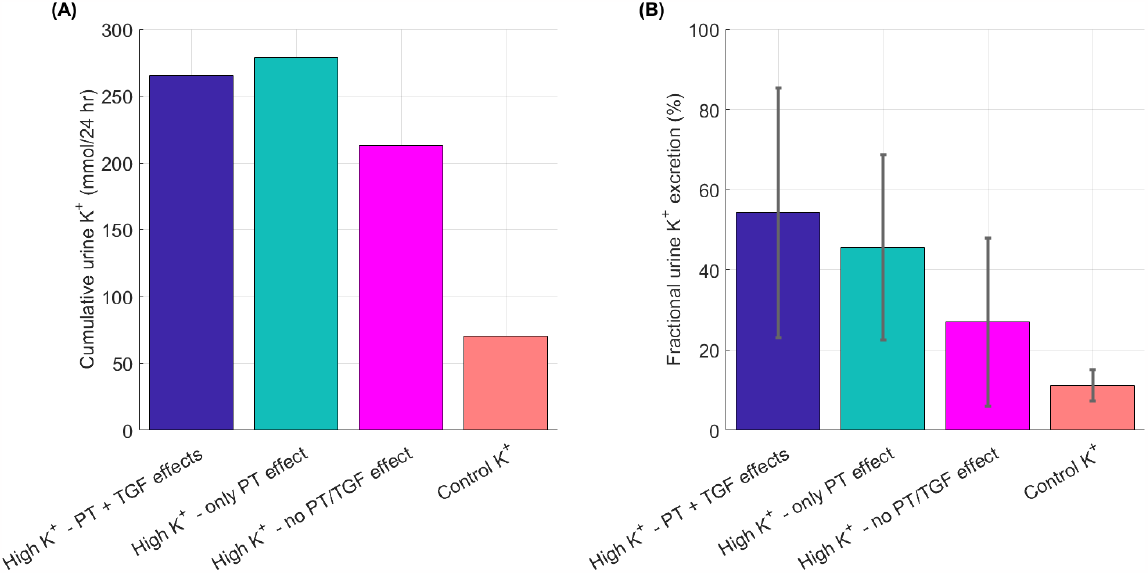
(A) Mean cumulative urine K^+^ excretion per day and (B) mean and standard deviation fractional urine K^+^ excretion over the full simulation time for control and high K^+^ diet with both PT and TGF effects (“PT + TGF effects”), only the PT effect (“only PT effect”), and no PT or TGF effects (“no PT/TGF effect”). PT: proximal tubule; TGF: tubuloglomerular feedback

**Figure 3.2:**
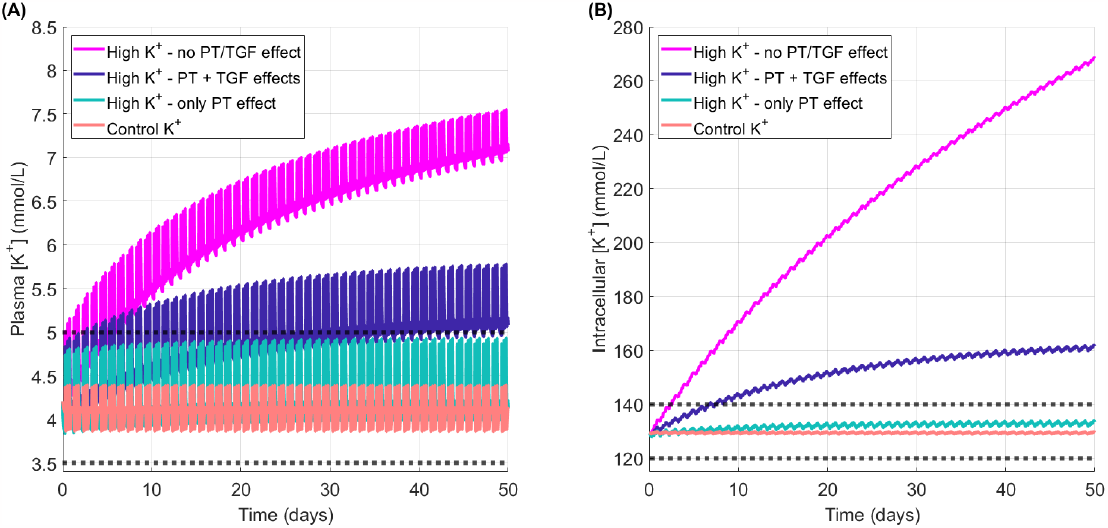
Predicted (A) plasma [K^+^] and (B) intracellular [K^+^] for control and high K^+^ diet with no PT or TGF effects, both PT and TGF effects, and only PT effect. Dark gray dotted lines indicate normal ranges for plasma [K^+^] (3.5-5.0 mmol/L) and intracellular [K^+^] (120-140 mmol/L). PT: proximal tubule; TGF: tubuloglomerular feeback

**Figure 3.3:**
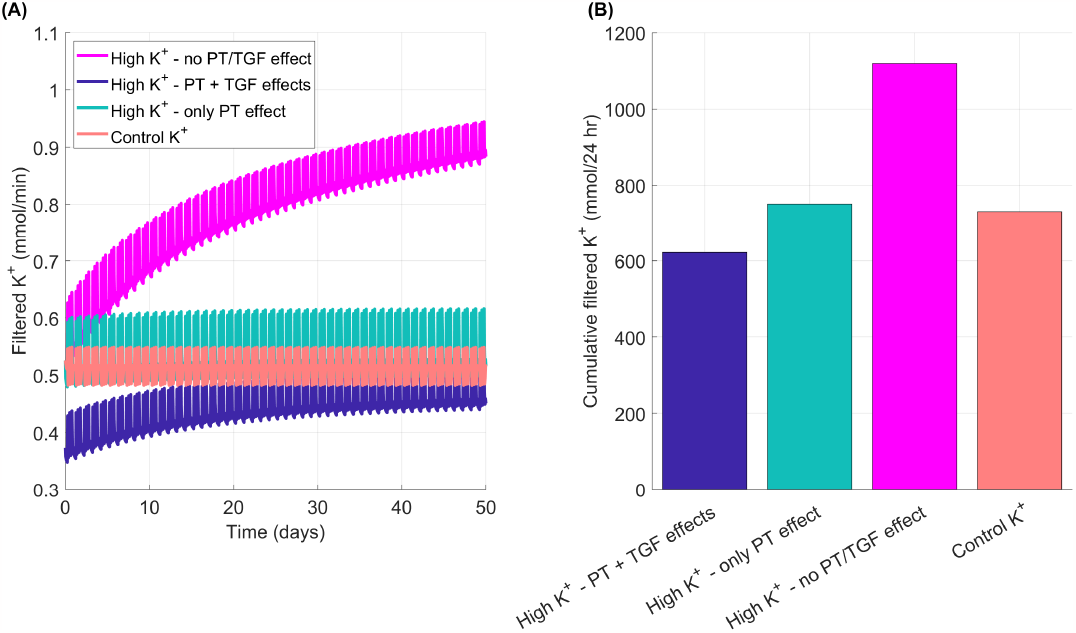
Filtered K^+^ into the nephrons (A) over the full simulation time and (B) mean cumulative filtered K^+^ per day. Simulation types correspond to Fig. 3.2.

**Figure 3.4:**
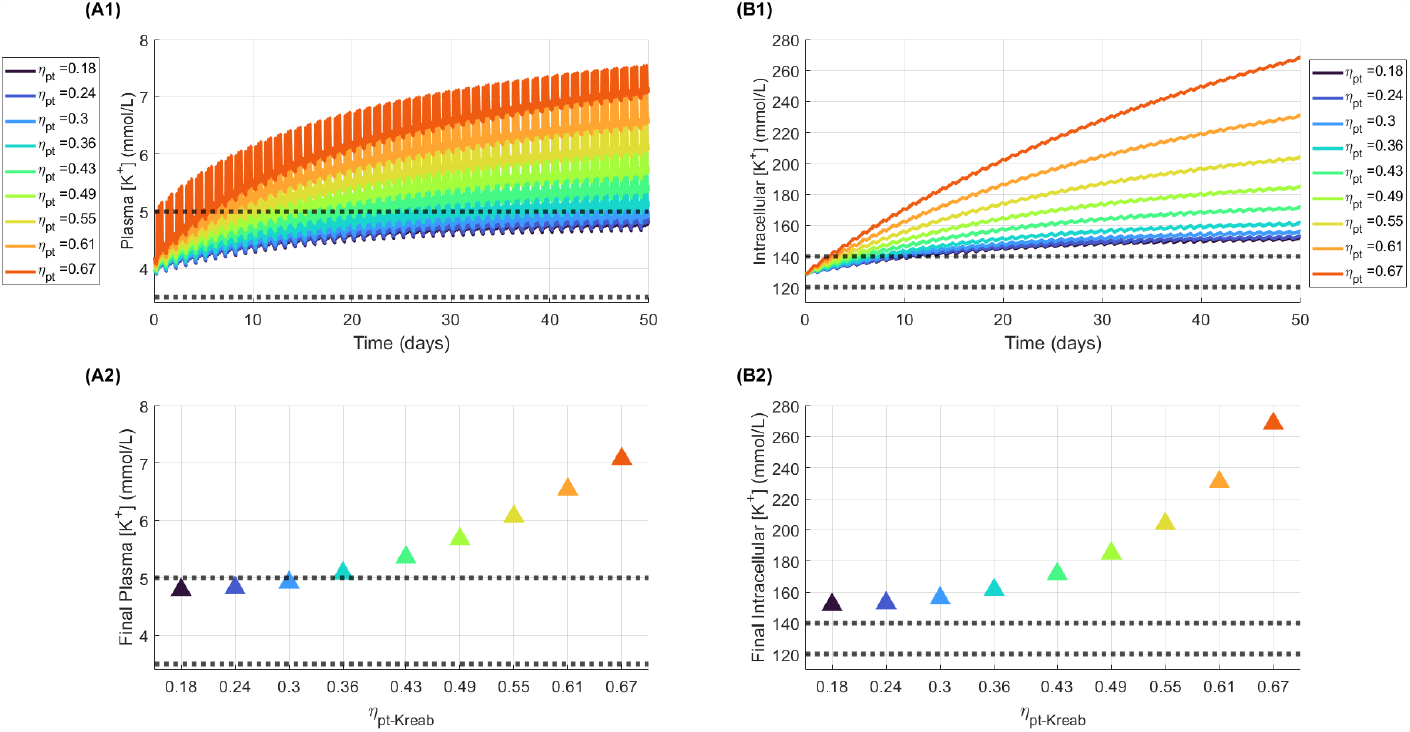
Predicted impact of varying fractional proximal tubule K^+^ reabsorption (η_pt−Kreab_, shortened to η_pt_ in the legends). Model predictions of plasma [K^+^] (A1) and intracellular [K^+^] (B1) for 50 day of high K^+^ diet with varied values for the η_pt−Kreab_ parameter, which represents fractional proximal tubule K^+^ reabsorption. The plasma and intracellular [K^+^] at the end of the 50 day simulations are plotted in panels A2 and B2, respectively. In all simulations shown here the TGF effect is included with 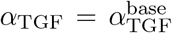. Note that the baseline high K^+^ intake fractional proximal tubule reabsorption is 0.36 as found by Wang et al. [42] and baseline normal K^+^ intake fractional proximal tubule reabsorption is 0.67. η_pt_: η_pt−Kreab_

**Figure 3.5:**
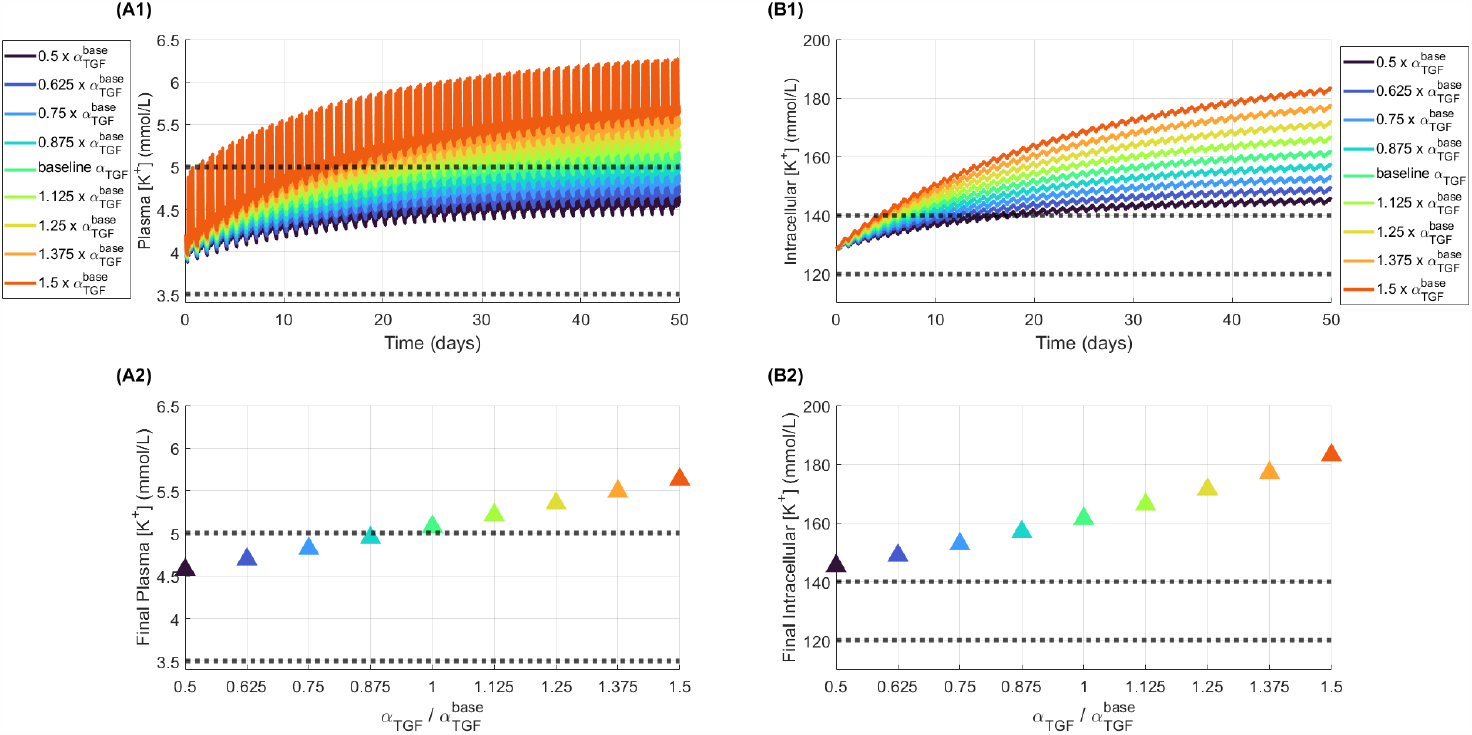
Predicted effects of varying the tubululoglomerular feedback (TGF) effect (α_TGF_). Model predictions of plasma [K^+^] (A1) and intracellular [K^+^] (B1) for 50 day of high K^+^ diet with varied values for the α_TGF_ parameter, which represents the strength of the TGF effect. The plasma and intracellular [K^+^] at the end of the 50 day simulations are plotted in panels A2 and B2, respectively. Proximal tubule K^+^ reabsorption η_pt−Kreab_ = 0.36 in all cases. 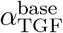 : baseline α_TGF_ parameter value (see Eq. 2.3)

**Figure 3.6:**
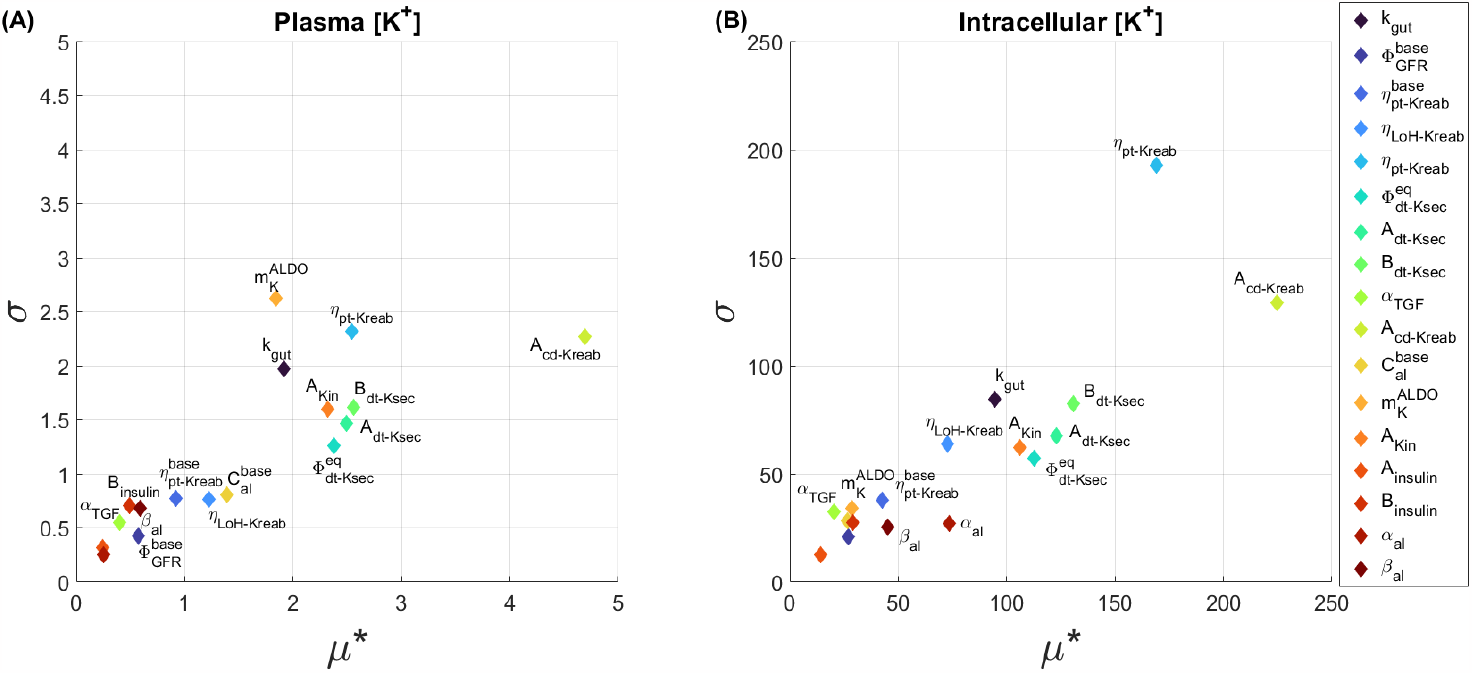
Morris plots for μ^∗^ (Eq. 2.8) and σ (Eq. 2.9) values for individual parameters values in the Morris analysis as measured by impact on the final (A) plasma [K^+^] and (B) intracellular [K^+^] at the end of a 50 day simulation of high K^+^ intake.

**Figure 3.7:**
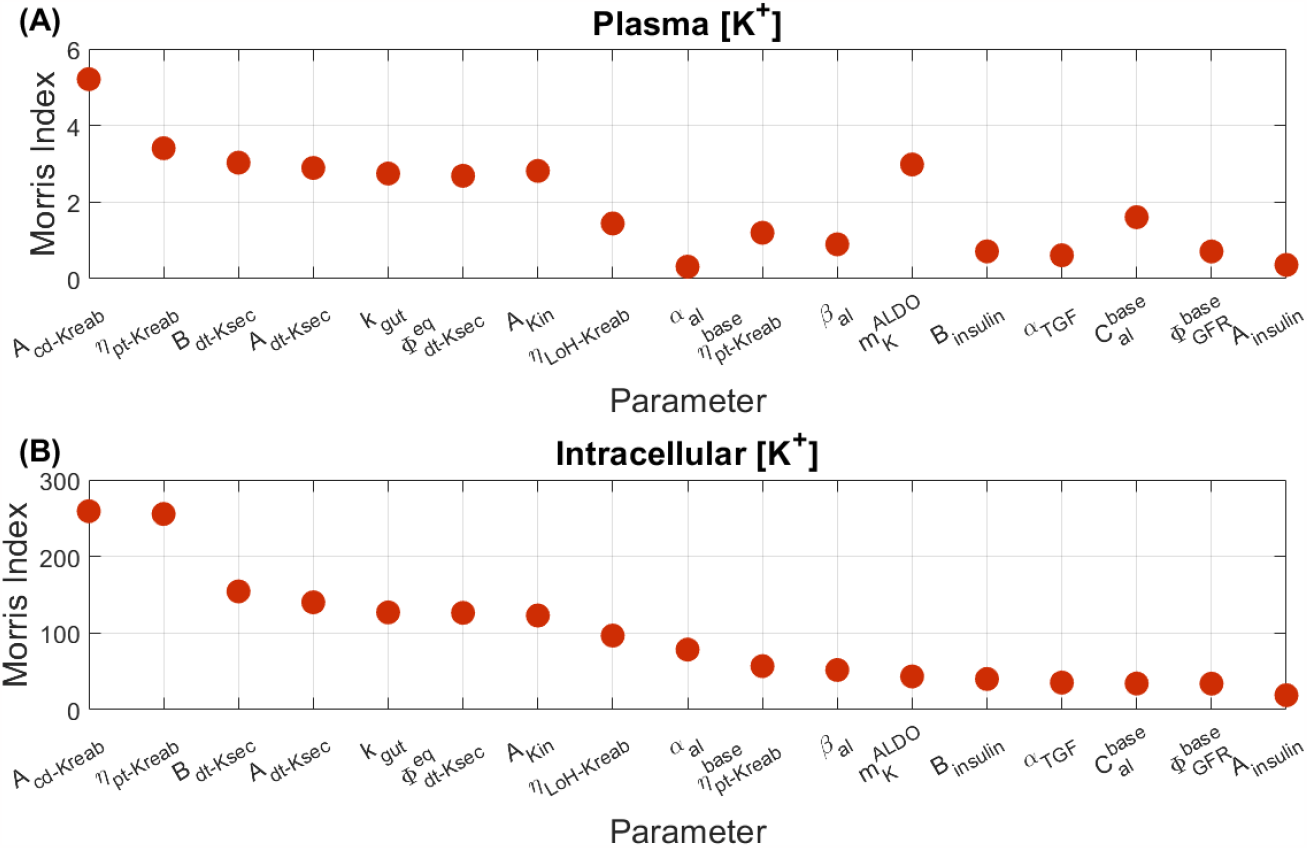
Morris indices (MI_i_; Eq. 2.10) for individual parameter values in the Morris analysis as measured by the impact on the final (A) plasma [K^+^] and (B) intracellular [K^+^] at the end of a 50 day simulation of high K^+^ intake.

**Figure 3.8:**
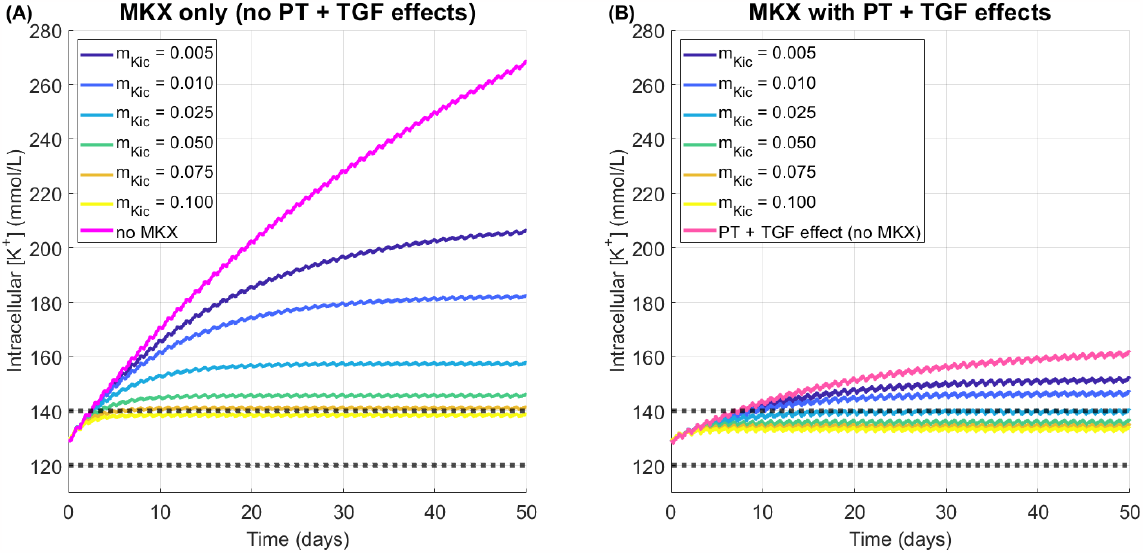
Model predictions of a high K^+^ diet for 50 days with and without muscle-kidney cross talk (MKX) and proximal tubule and tubuloglomerular feedback effects (PT + TGF). PT + TGF effects includes Eq. 2.2 with η_pt−Kreab_ = 0.36 as in Wang et al. [42] and α_TGF_ is the baseline value as given by Eq. 2.3. (A) Model predictions of intracellular [K^+^] with muscle-kidney cross talk only where parameter m_Kic_ is the value as given by legend. (B) Model predictions of intracellular [K^+^] with muscle-kidney cross talk with PT + GFR high K^+^ effects where parameter m_Kic_ is as given by the legend. Normal range for intracellular [K^+^] indicated by gray dotted line (120-140 mmol/L). MKX: muscle-kidney cross talk; PT: proximal tubule; TGF: tubuloglomerular feedback; m_Kic_: muscle-kidney cross talk parameter (see Eq. 2.11).

**Figure 3.9:**
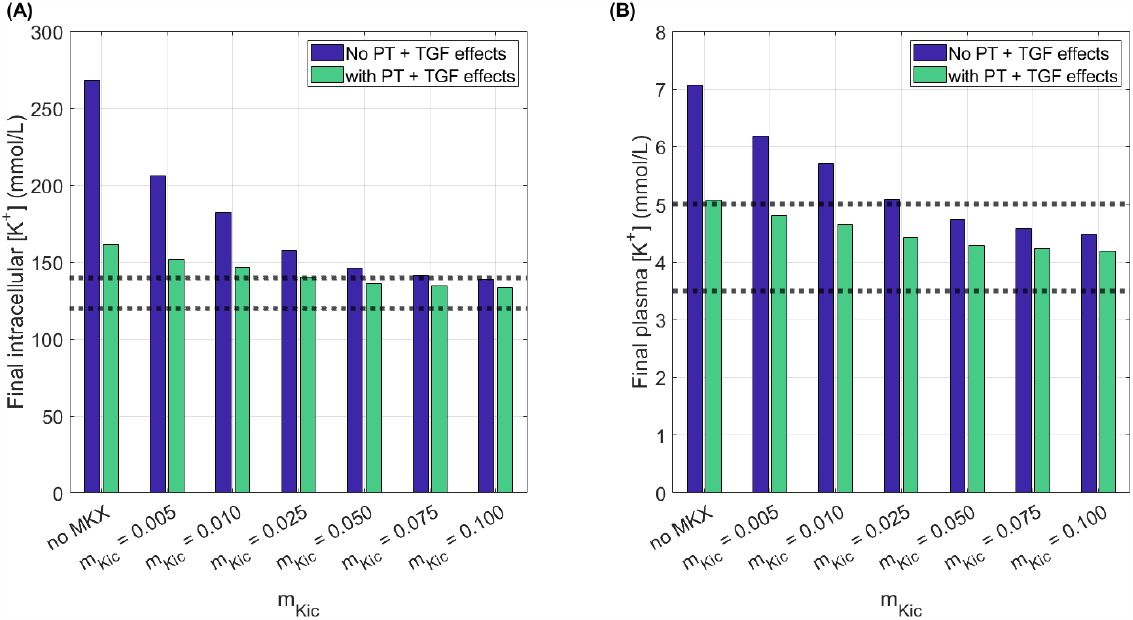
(A) Intracellular and (B) plasma [K^+^] levels at the end of 50 day simulation under with and without proximal tubule and tubuloglomerular feedback effects (PT + TGF effects). Muscle-kidney cross talk as given by the parameter m_Kic_ with or without PT + GFR effects. Normal ranges for intracellular and plasma [K^+^] indicated by gray dotted line (120-140 mmol/L and 3.5-5.0 mmol/L, respectively). PT: proximal tubule; TGF: tubuloglomerular feedback; m_Kic_: muscle-kidney cross talk parameter (see Eq. 2.11).

For the high K^+^ diet simulation, total daily K^+^ intake is 312 mmol, 10% of which is assumed to be excreted via feces (or other mechanisms such as sweat). To achieve K^+^ balance, the remaining 281 mmol should be excreted via urine. Without PT K^+^ reabsorption adaptation to a high K^+^ diet (“no PT/TGF effect”), daily urinary K^+^ excretion is about 213 mmol, resulting in a 68 mmol per day deficit for K^+^ balance (Fig. 3.1A). Predicted fractional urinary K^+^ excretion for the simulation of a high K^+^ diet with no PT or TGF effects is on average 27% (Fig. 3.1B; “no PT/TGF effect”), below the 30%–130% range reported for rats on a high K^+^ diet [42]. The predicted lower fractional and daily urine K^+^ excretions results in an imbalance between K^+^ intake and excretion; as a result, total body K^+^ increases. By day 3 of the simulation time, predicted intracellular [K^+^] levels are above normal range (*>*140 mmol/L) and by day 5 of simulation time predicted plasma [K^+^] is above normal range (*>*5.0 mmol/L) (Fig. 3.2; “no PT/TGF effect”). As the simulation time continues, plasma and intracellular [K^+^] levels reach dangerously high levels at over 7 mmol/L and 260 mmol/L, respectively.

With the addition of the PT and TGF effects, predicted fractional urinary K^+^ excretion is on average 54% (Fig. 3.1B; “PT + TGF effects”), within the range found by Wang et al. [42]. This results in daily cumulative urine K^+^ excretion of 265 mmol (Fig. 3.1A; “PT + TGF effects”), only 5% below the excretion needed for perfect balance with K^+^ intake. Predicted plasma and intracellular [K^+^] levels do increase as a result of this imbalance, but at a much slower rate than when there are no PT and TGF effects (Fig. 3.2; “PT + TGF effects”). By day 10, intracellular [K^+^] is above normal range and plasma [K^+^] is consistently above normal range by day 20 (Fig. 3.2). At the end of the simulation, plasma [K^+^] is 5.1 mmol/L and intracellular [K^+^] is 162 mmol/L (Fig. 3.2).

With a high K^+^ diet, the reduction in PT K^+^ reabsorption increases K^+^ excretion, whereas TGF lowers GFR and, when taken in isolation, reduces K^+^ excretion. To uncouple the effects of these two opposing responses, we consider the hypothetical situation with only the reduction in PT K^+^ reabsorption, but no TGF effect, i.e., GFR is kept at baseline level. The model simulation predicts that daily cumulative urine K^+^ excretion is nearly at exact balance at 279 mmol – 90% of K^+^ intake of 312 mmol is 280.8 mmol (Fig. 3.1A; “only PT effect”). This results in nearly steady intracellular [K^+^] levels (Fig. 3.2B; “only PT effect”). At the end of the simulation time, plasma [K^+^] is 4.2 mmol/L and intracellular [K^+^] is 133.4 mmol/L, both well within normal range (Fig. 3.2). Note that while removing the TGF increases net urinary K^+^ excretion, it also decreases fractional urine K^+^ excretion (Fig. 3.1B; “only PT effect”). This apparent paradox is due to the GFR-lowering effect of TGF (Eq. 2.2), which decreases K^+^ filtration. A further analysis of the dynamics of filtered K^+^ revealed that, in the first 10 days of the simulation, filtered K^+^ is on average 24% lower with the TGF effect (compared “PT + TGF effects” with “only PT effect” high K^+^ simulation in Fig. 3.3A). With the TGF effect represented, filtered K^+^ increases over time due to the increasing plasma [K^+^] levels (Fig. 3.3A; Fig. 3.2A). Despite that increase, average cumulative filtered K^+^ per day is 17% lower for the “PT + TGF effects” simulation than the “only PT effect” simulation over the 50 days (Fig. 3.3B).

In summary, adaptive PT K^+^ reabsorption to a high K^+^ diet found in Ref. [42] is sufficient for maintaining plasma and intracellular [K^+^] levels within normal range, if it were the only mechanism activated. However, the TGF effect caused by altered NaCl and K^+^ delivery to the macula densa reduces K^+^ filtration. This results in lower urinary K^+^ excretion causing significant K^+^ imbalance. Ultimately, the K^+^ retention leads to a high total body K^+^ content as shown by increased plasma and intracellular [K^+^] levels on a high K^+^ diet.

### 3.2 Predicted effects of varying fractional proximal tubule K^+^ reabsorption and tubuloglomerular feedback

The extent to which PT K^+^ reabsorption is suppressed under a high K^+^ diet may vary due to factors such as Na^+^ intake, hormone levels, and other physiological conditions. How does variations in PT K^+^ reabsorption impact whole body K^+^ handling under a high K^+^ diet? To investigate this question we conducted high K^+^ intake simulations for varied values of fractional PT K^+^ reabsorption (*η*_pt−Kreab_; Eq. 2.1). These simulations were done with the TGF effect. Results are shown in Fig. 3.4.

Recall that, for no change in *η*_pt−Kreab_ under high K^+^ intake (*η*_pt−Kreab_ = 0.67), plasma and intracellular [K^+^] quickly exceed normal range (Fig. 3.2; Fig. 3.4A1-B1). Decreasing *η*_pt−Kreab_ by 0.06 from baseline leads to a 7.5 and 14% decrease in plasma and intracellular [K^+^] at the end of the simulation, respectively (Fig. 3.4A2-B2 *η*_pt−Kreab_ = 0.61). A similar decrease in plasma and intracellular [K^+^] is predicted for another decrease by 0.06 (Fig. 3.4A2-B2, *η*_pt−Kreab_ = 0.55). However, as we continue to decrease *η*_pt−Kreab_ by 0.06, the model predicts diminishing responses, such that when decreasing *η*_pt−Kreab_ from 0.24 to 0.18, there is a negligible difference in the end point plasma and intracellular [K^+^] (Fig. 3.4A2-B2). This is due to the TGF signal, which decreases GFR when delivery of K^+^ to the macula densa is lowered as a result of decreased PT K^+^ reabsorption.

The TGF signal can vary due to factors other than PT K^+^ reabsorption (e.g., Na^+^ or Cl^−^ levels). To investigate the impact of a differing TGF signal, we varied the parameter *α*_TGF_ from 0.5 to 1.5 times the baseline value 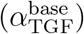 and conducted simulations for a high K^+^ diet with *η*_pt−Kreab_ = 0.36 (fractional PT K^+^ reabsorption under a high K^+^ diet as reported in Ref. [42]). The results are shown in Fig. 3.5.

Note that a stronger TGF signal (larger *α*_TGF_) results in a lower GFR based on the lower *η*_ps−Kreab_ under a high K^+^ diet (Eq. 2.2). This results in a faster increase in plasma and intracellular [K^+^] levels (Fig. 3.5A1-B1). Conversely, when decreasing *α*_TGF_, the GFR is decreased less by the reduced PT K^+^ reabsorption, resulting in a slower increase in plasma and intracellular [K^+^] levels due to higher K^+^ filtration than 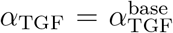 (Fig. 3.5A1-B1). When varying *α*_TGF_, there is a linear response in the final plasma and intracellular [K^+^] levels (Fig. 3.5A2-B2). For 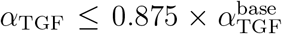, the plasma [K^+^] stays within normal range for the full simulation (Fig. 3.5A1-A2). However, for all *α*_TGF_ values simulated, intracellular [K^+^] is higher than its normal range (Fig. 3.5B1).

In summary, decreasing fractional PT K^+^ reabsorption under a high K^+^ diet improves plasma and intracellular [K^+^] levels in a nonlinear trend so that there is a limiting effect on the impact of lowering fractional PT K^+^ reabsorption. This nonlinearity is due to the TGF effect, which lowers GFR in response to high delivery of K^+^ to the macula densa. Similarly, decreasing the TGF effect parameter, *α*_TGF_ results in a linear improvement in plasma and intracellular [K^+^]. However, the presence of a TGF signal, even a weak one, would eventually yield intracellular [K^+^] that exceeds its normal range. In both cases, plasma [K^+^] remained in normal range for sufficiently low fractional PT K^+^ reabsorption and *α*_TGF_ values.

### 3.3 Sensitivity analysis

Regulation of K^+^ can be impacted by many different physiological changes, represented in the model by varying parameter values. In order to analyze the extent that variations in different parameters impact final plasma and intracellular [K^+^], we conducted a Morris analysis of key parameters.

The parameters included in the Morris analysis were determined by a local sensitivity analysis to reduce computation time. The results for parameters that had a more than 2% change in *K*_plasma_ or *K*_intracellular_ in the local sensitivity analysis (Eq. 2.6) are shown in Fig. A.2 and are the ones that are included in the Morris analysis. See Table B.1 for the parameter ranges used for the Morris analysis. There are 17 parameters that were included in the Morris analysis.

Figure 3.6 shows the *μ*^∗^ and *σ* values (Eq. 2.8; Eq. 2.9) for each of the parameters included in the Morris analysis with respect to the final plasma and intracellular [K^+^]. This type of plot is often referred to as a “Morris plot” [31]. The Morris indices (Eq. 2.10) are shown in Fig. 3.7. Any parameters not included had an inconsequential impact on final plasma and intracellular [K^+^] levels.

The *μ*^∗^ and *σ* values for the intracellular [K^+^] are much larger than the plasma [K^+^] (Fig. 3.6). This is due to intracellular [K^+^] having a larger baseline value than plasma [K^+^] (130 mmol/L vs. 4.2 mmol/L) and intracellular [K^+^] having a larger tolerance for changes in K^+^ levels. Hence, it is expected that there should be more variance in the final intracellular [K^+^] than plasma [K^+^] as reflected in the Morris analysis results.

The parameters with the three largest Morris indices for both plasma and intracellular [K^+^] are *A*_cd−Kreab_, *η*_pt−Kreab_, and *B*_dt−Ksec_ (Fig. 3.7), which impact collecting duct K^+^ reabsorption, PT K^+^ reabsorption, and distal tubule K^+^ secretion, respectably. These are all parameters that are involved in renal handling of K^+^.

The parameter with the fourth largest Morris index for plasma [K^+^] is 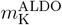 (Fig. 3.7A), which determines how changes in the extracellular [K^+^] regulates ALD secretion. This parameter has a relatively large Morris index for the plasma [K^+^] because of its large value for *σ* (Fig. 3.6A), therefore suggesting that this parameter likely interacts highly with other parameters. In the local sensitivity analysis, 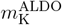 did not have a high impact.

In this study, the primary model extension was the renal effects on a high K^+^ diet, by changing *η*_pt−Kreab_ and adding Eq. 2.2 to include TGF effects. Indeed, in the Morris analysis we found that both plasma and intracellular [K^+^] are highly sensitive to changes in *η*_pt−Kreab_ (Fig. 3.6; Fig. 3.7) on a high K^+^ diet. In contrast, *α*_TGF_ had a smaller impact on the model output (Fig. 3.6; Fig. 3.7). Note that 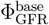, which represents the baseline GFR, also has a relatively low Morris index for both plasma and intracellular [K^+^] (Fig. 3.7). These results suggest that the final plasma and intracellular [K^+^] are affected more by changes in K^+^ transport along the nephron segments than by filtration.

In summary, our Morris analysis results quantify the impacts of each parameter and its interactions with other parameters on the final plasma and intracellular [K^+^] after 50 days of high K^+^ intake. Our Morris analysis results show that final plasma and intracellular [K^+^] levels depend primarily on parameters that characterize renal K^+^ transport.

### 3.4 Muscle-kidney cross talk simulations

To investigate the impact of the hypothesized muscle-kidney cross talk signal in our updated mathematical model, we conduct “what-if” simulations of a high K^+^ diet and vary the muscle-kidney cross signal “strength,” denoted by *m*_Kic_ (Eq. 2.11).

Results for predicted intracellular [K^+^] during high K^+^ diet simulations of muscle-kidney cross talk for varied *m*_Kic_ values with the PT + TGF effects are shown in Fig. 3.8A. A stronger cross talk signal, i.e., larger *m*_Kic_, results in lower intracellular [K^+^] levels. When *m*_Kic_ = 0.1, intracellular [K^+^] levels remain within normal range for the full simulation. However, the predicted intracellular [K^+^] are on the high end of the normal range. Results for the same simulations with the addition of PT + TGF effects (*η*_pt−Kreab_ = 0.36 and 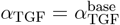) are shown in Fig. 3.8B. We can see that with the PT + TGF effects and muscle-kidney cross talk parameter *m*_Kic_ ≥ 0.025, intracellular [K^+^] levels remain within the normal range for the full simulation (Fig. 3.8B).

The intracellular and plasma [K^+^] at the end of the 50 day high K^+^ simulations with and without PT + TGF effects for varied *m*_Kic_ are shown in Fig. 3.9. Regardless of PT + TGF effects, increasing *m*_Kic_ decreases both final intracellular and plasma [K^+^] levels (Fig. 3.9). As expected, adding the PT + TGF effects improved final intracellular and plasma [K^+^] levels. This is especially true for the no muscle-kidney cross talk simulation or small values of *m*_Kic_. Final plasma [K^+^] levels are within normal range with the addition of the muscle-kidney cross talk for *m*_Kic_ ≥ 0.005 (Fig. 3.9B), but final intracellular [K^+^] remains higher than normal range until *m*_Kic_ ≥ 0.025 (Fig. 3.9A).

In summary, while the impacts of high K^+^ intake on PT reabsorption greatly improve the model’s plasma and intracellular [K^+^] predictions, the presence of a muscle-kidney cross talk signal prevents excess K^+^ from being stored in the intracellular fluid.

## 4 Discussion

The mathematical model presented in this study includes a comprehensive description of known regulatory mechanisms involved in K^+^ handling, as well as the newly discovered impacts of high K^+^ intake on PT K^+^ reabsorption and the resulting TGF effects [42]. We conducted model analysis to:

- Predict the impact of PT and TGF effects of a high K^+^ diet on whole body potassium homeostasis (Fig. 3.1; Fig. 3.2; Fig. 3.3).
- Unravel the interconnected effects of decreased PT K^+^ reabsorption with the TGF signal under a high K^+^ diet (Fig. 3.1; Fig. 3.2; Fig. 3.3).
- Quantify the impact of varied amounts of fractional PT K^+^ reabsorption (*η*_pt−Kreab_; Fig. 3.4) and TGF signaling (*α*_TGF_; Fig. 3.5) under a high K^+^ diet.
- Determine which parameters have the largest impact on the final plasma and intracellular [K^+^] under high K^+^ intake from a global approach (Fig. 3.6; Fig. 3.7).
- Investigate the hypothesized muscle-kidney cross talk signal under a high K^+^ diet with PT + TGF effects via “what-if” simulations (Fig. 3.8; Fig. 3.9).

Overall, our model analysis provides insights into the impacts of renal adaptations to a modified K^+^ diet, and provides a better understanding of the complex regulation of K^+^ homeostasis under high K^+^ intake.

The kidneys play a key role in long-term K^+^ homeostasis by ensuring that total daily K^+^ excretion matches daily K^+^ intake. Detailed mathematical models of renal epithelial transport have been developed and applied to K^+^ handling by the kidney, albeit without explicitly considering extrarenal K^+^ balance. The model of rat kidney function by Weinstein [43] indicated that, a K^+^ load attenuates PT fluid reabsorption and increases Na^+^ delivery to the distal nephron, despite an TGF-lowered GFR. As previously reported in Ref. [16], our group’s model simulations of nephron function following an acute K^+^ load indicated that elevated expression levels and activities of Na^+^/K^+^-ATPase, epithelial sodium channels, large-conductance Ca^2+^-activated K^+^ channels, and renal outer medullary K^+^ channels, together with downregulation of Na^+^-Cl^−^ cotransporters, increase K^+^ secretion along the connecting tubule, resulting in a *>*6-fold increase in urinary K^+^ excretion, which substantially exceeds the filtered K^+^ load.

Unlike the studies discussed above, the kidney component of the present model is much simpler and does not represent detailed epithelial transport [16, 43]. Without altering nephron transport under a high K^+^ diet, urinary K^+^ excretion is too low to maintain K^+^ balance (Fig. 3.1A). By reducing the fractional PT K^+^ reabsorption, average fractional urine K^+^ excretion nearly doubles, resulting in near K^+^ balance (Fig. 3.1B). However, changes to electrolyte transport in the kidneys does not occur in isolation. Notably, the decrease in PT K^+^ reabsorption found by Wang et al. [42] is driven by reduced Na^+^ reabsorption by Na^+^ transporters. That results in a higher Na^+^ and K^+^ load to the macula densa, where Na^+^, K^+^, and Cl^−^ levels are detected for TGF signalling. The TGF signal reduces flow to the glomerulus, lowering the single nephron GFR, and therefore directly decreasing renal K^+^ filtration into each nephron. The resulting reduction in urinary K^+^ excretion leads to a slight K^+^ imbalance (Fig. 3.1A) and a gradual rise in total body K^+^ levels (Fig. 3.2). Our model simulations unravel the complex interplay of K^+^ transport along the PT with TGF effects.

Heterogeneity in individual biology impacts how the body responds to different physiological stimuli. To explore the functional implications of individual differences, we conducted simulations with different decreases in PT K^+^ reabsorption under a high K^+^ diet by changing the relevant parameter value (*η*_pt−Kreab_; Eq. 2.1). We found that due to the TGF signal, decreasing PT K^+^ reabsorption has a diminishing impact on plasma and intracellular [K^+^] balance (Fig. 3.4). Specifically, we simulated changes in *η*_pt−Kreab_ to as low as 50% of the reported fractional PT K^+^ reabsorption in Ref. [42] and found that at the end of the 50 day simulation, plasma [K^+^] was on the high end of normal range in between meals and intracellular [K^+^] were above normal range by day 20 (Fig. 3.4). While in this case plasma [K^+^] is within the normal range, a person with high intracellular [K^+^] levels may be more susceptible to hyperkalemia because of impaired intracellular K^+^ buffering. Additionally, we note that this low fractional PT K^+^ reabsorption may be outside of physiological range.

Heterogeneity in physiology also impacts the strength of the TGF response. To investigate this, we conducted high K^+^ intake simulations with varied TGF strength characterized by *α*_TGF_ (Eq. 2.2). When *α*_TGF_ is increased, the plasma and intracellular [K^+^] levels increase as a result of decreased filtration to the nephrons (Fig. 3.5). Conversely, when *α*_TGF_ is decreased, the filtration is not as strongly affected by reduced PT fractional K^+^ reabsorption, which yields a better balance of urinary K^+^ excretion. When *α*_TGF_ is reduced by 12.5% or more, final plasma [K^+^] is within normal range (Fig. 3.5A2), but even when *α*_TGF_ is half of its baseline value (i.e., 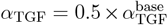), intracellular [K^+^] levels rise to higher than normal range within the 50 day simulation (Fig. 3.5B1-B2). This is because even with a smaller reduction in K^+^ filtration, the change is enough to result in deficit urinary K^+^ excretion so that some K^+^ is retained within the body. Since the intracellular fluid uptakes extra K^+^ to keep plasma [K^+^] within normal range, intracellular [K^+^] levels are increased. Therefore, the TGF effect lowers K^+^ filtration leading to reduced urinary K^+^ excretion so that excess K^+^ remains within the body.

As with many mathematical models of physiological systems, the present model contains (i) epistemic uncertainty, which comes from an incomplete knowledge of the physiological system, and (ii) aleatory uncertainty, which refers to randomness and heterogeneity that arise in biological systems [34]. For example, in humans, it is impossible to (ethically) measure the exact rate of how K^+^ secretion along the distal tubule changes with changes in [ALD], rather these parameters must be estimated based on the data that we do have. This is an example of epistemic uncertainty in the K^+^ regulation model. Aleatory uncertainty comes in that even if we could measure every parameter exactly in a person, this will not represent the whole population, and may not even represent that person on a different diet or at a different time. For this reason, studying the effects of varied parameters using global sensitivity analysis can help us further understand uncertainties that arise when modeling complex systems as done in this study.

To elucidate the relationship of the parameters and the predicted plasma and intracellular [K^+^], we conducted a Morris analysis to quantify how much each parameter impacts final plasma and intracellular [K^+^] after 50 days of a high K^+^ diet (Fig. 3.6; Fig. 3.7). This analysis showed that for both plasma and intracellular [K^+^], the parameters that determine renal K^+^ transport have the largest impact on final [K^+^]. In particular, the parameter with the second highest impact was the PT K^+^ reabsorption, showing that this change under a high K^+^ diet is critical for maintaining K^+^ balance even when other parameters are altered. Additionally, this Morris analysis showed that our model predictions were driven primarily by renal transport. This shows that our model captures what is happening physiologically since long-term K^+^ regulation is primarily regulated by renal K^+^ handling via urinary K^+^ excretion.

In our previous study [39], we used our mathematical model to conduct “what-if” simulations of muscle-kidney cross talk under short-term K^+^ loading or depletion. Specifically, we found that in our baseline model (i.e., no muscle-kidney cross talk) after 4 days of K^+^ loading (high K^+^ intake), plasma [K^+^] levels started to reach hyperkalemic levels and intracellular [K^+^] levels remained high for several days after returning to normal K^+^ intake [39]. When the muscle-kidney cross talk signal was included, model simulations predicted that intracellular [K^+^] levels remained within normal range. In this study, we extended our model to consider PT and TGF effects. Without the muscle-kidney cross talk signal, these effects were unable to keep both plasma and intracellular [K^+^] levels within normal ranges on a chronic high K^+^ diet (Fig. 3.2), which suggests that the muscle-kidney cross talk signal may be one of the necessary regulatory mechanisms. To test this, we conducted “what-if” simulations of a high K^+^ diet with varying strengths of the muscle-kidney cross talk signal (*m*_Kic_; Eq. 2.11). Model results indicated that muscle-kidney cross talk significantly improved K^+^ balance (Fig. 3.8; Fig. 3.9).

The mechanisms that regulate K^+^ balance are highly coupled. Mathematical modeling is a useful tool for unraveling complex systems by capturing the highly coupled nature and using simulations and analysis to investigate the intricacies of the system. One advantage is that we can conduct hypothetical experiments that would not be possible *in vivo* such as predicting whole-body K^+^ homeostasis with varied PT K^+^ reabsorption and TGF effects as we did in this study. This analysis can be used to understand the system and make hypotheses to guide future work. In this study, we extended our model to capture renal effects under a high K^+^ diet. This is an important step for understanding how changes in organs such as the kidneys impact whole-body K^+^ homeostasis. Our model and analysis provide a framework for further studies of K^+^ homeostasis in health and disease, as well as for the development of new strategies to prevent and treat K^+^-related disorders.

### Model limitations and future work

Since this study is an extension of our previously built model (Ref. [39]), similar limitations apply here. First of all, we note that regulation of K^+^ transport as well as the TGF signal is indeed coupled to other solutes such as Na^+^. While we do not include Na^+^ regulation in this study explicitly, indeed this model could be combined with an analogous model of Na^+^ regulation (e.g., Refs. [1,7,19]). The resulting model can be used to investigate the synergy of Na^+^ and K^+^ balance in blood pressure control.

We note that the whole-body K^+^ regulation model is for a human (man). However, the addition of the PT K^+^ reabsorption adaptations under a high K^+^ diet is from a rat study [42]. Indeed, there are species differences in renal handling [18, 38]. However, due to the invasive experiments required to get renal transport data as done in Ref. [42], human data is not possible at this time. Therefore, we assumed that the same percent change in fractional PT K^+^ reabsorption and TGF effects occured in the human model, but did not change other renal parameters. It is likely that there may be variation between the species, which is why studying differences in paramters such as done in Fig. 3.4 and Fig. 3.5 is useful for studying heterogeneity that may occur in populations and between the species.

Another limitation and opportunity is that this model is built on data that is not sex-specific. It is known that there are significant sexual dimorphisms in renal K^+^ transport [10,24,38], Na^+^ balance [24, 38], and ALD concentration [26], which all affect K^+^ homeostasis. Sex-specific K^+^ models will need to be developed to capture female-specific K^+^ regulation. It has also been hypothesized that males and females have different plasma [K^+^] set points than males for handling high K^+^ loads [28]. Indeed, this whole-body K^+^ model may be extended to consider sex differences as well as unique physiological states in females that undergo major changes in K^+^ regulation such as pregnancy and lactation [37, 40, 45].

Recent work has revealed the impacts of circadian rhythms on renal electrolyte handling, particularly changes in K^+^ transport [3, 6, 17, 36]. Indeed, this likely has a role in whole-body K^+^ homeostasis. A similar approach to this current study may include circadian effects on renal K^+^ transport to investigate how this impacts the full body.

Understanding K^+^ homeostasis is critical due to the risks and potentially severe consequences of hyper- and hypokalemia. Our current study considers K^+^ regulation in a healthy person. Indeed, future work may consider pathological states such as chronic kidney disease or persons taking medications that can affect K^+^ balance. To simulate impaired kidney function or medications that target the kidney, the renal component of the present model may be replaced by one that represents detailed epithelial transport (e.g., [9, 41]).

## A Additional Figures

### A.1 High K^+^ intake

**Figure A.1:**
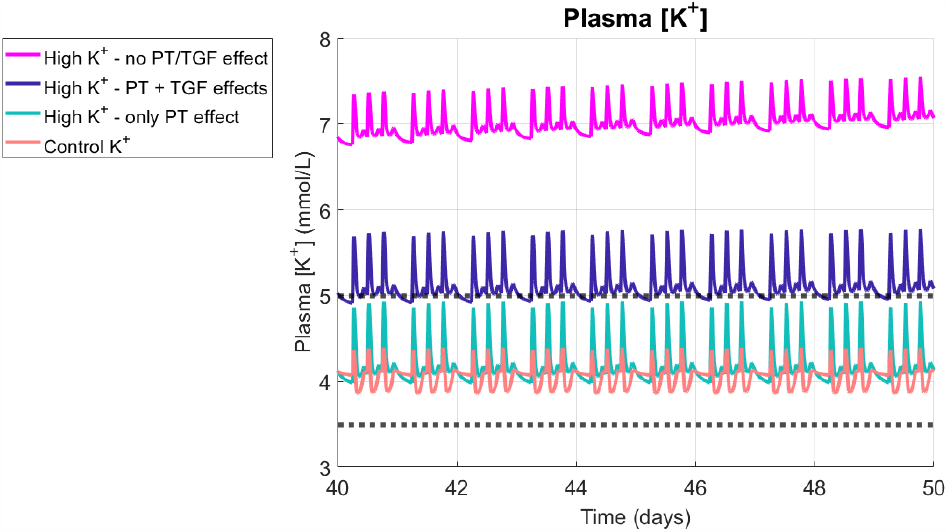
Replication of Fig. 3.2A for the last 10 days of the model simulation.

### A.2 Sensitivity analysis

**Figure A.2:**
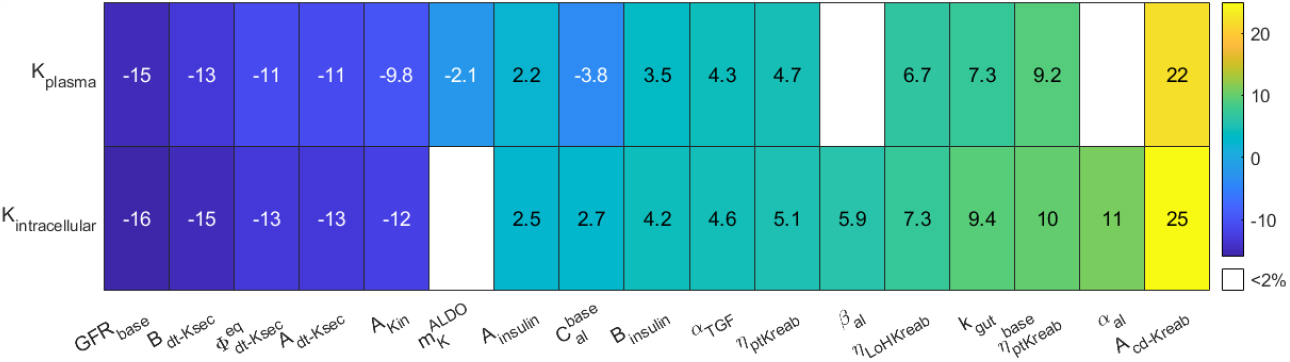
Local sensitivity analysis of impact on individual parameters on the end plasma and intracellular [K^+^] at the end of the 50 day simulation. Sensitivity is determined by increasing and decreasing individual parameters by 10%.Parameters that did not result in a greater than 1% change for either K_plasma_ or K_intracellular_ are not shown. Sensitivity was determined by Eq. 2.6. K_plasma_: plasma [K^+^] at end of 50 day simulation; K_intracellular_: intracellular [K^+^] at end of 50 day simulation.

## B. Description of baseline mathematical model of potassium homeostasis regulation

Here we describe the mathematical model of potassium homeostasis used in our study. For more details see Stadt et al. [39]. A list of the parameters and their values used in the baseline model is given in Table B.1.

### B.1 Internal K^+^ balance

Internal K^+^ is divided into 4 compartments: gastrointestinal and hepatoportal circulation (denoted by *M*_Kgut_), plasma [K^+^] (denoted by *K*_plasma_), interstitial fluid [K^+^] (*K*_inter_), and intracellular [K^+^] (*K*_IC_).

The amount of K^+^ in the gut and hepatoportal circulation is given by

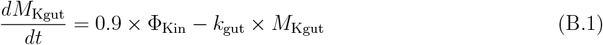

where Φ_Kin_ is K^+^ intake and *k*_gut_ is a parameter that determines how fast the K^+^ is delivered to the plasma. Plasma [K^+^] concentration is given by

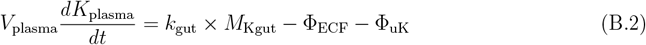

where *V*_plasma_ is plasma volume, Φ_uK_ denotes urine K^+^ excretion (described in Section B.2); Φ_ECF_ denotes the diffusion of K^+^ from the blood plasma to the interstitial space and is given by

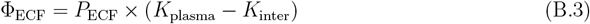

where *P*_ECF_ is a permeability parameter. Interstitial [K^+^] is determined by

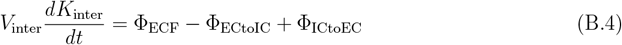

where *V*_inter_ is the interstitial fluid volume, Φ_ECtoIC_ and Φ_ICtoEC_ are the fluxes of K^+^ from the extracellular to the intracellular fluid and vice versa, respectively (see Eqs. B.6 & B.7). The intracellular [K^+^] is determined by the fluxes of K^+^ in and out of the cells

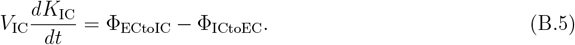

Flow of K^+^ into the intracellular fluid is driven by Na^+^-K^+^-ATPase uptake and is modeled using Michaelis-Menten kinetics so that

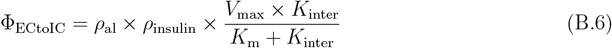

where *V*_max_ is the maximum rate, *K*_m_ denotes the half maximal activation level, *ρ*_al_ is the effect of ALD on Na^+^-K^+^-ATPase (see Section B.3), and *ρ*_insulin_ is the effect of insulin (see Section B.4). K^+^ returns to the extracellular compartment via diffusion through a permeable membrane:

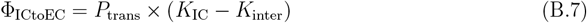

where *P*_trans_ denotes the transmembrane permeability.

### B.2 Renal K^+^ regulation

Filtered K^+^ load (Φ_filK_) is proportional to the GFR (Φ_GFR_) and plasma [K^+^] so that

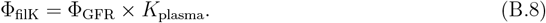

The filtrate then moves through the nephrons, where K^+^ is reabsorbed and secreted along the different segments. At the end of the nephrons, what is remaining is excreted in urine. The model represents the kidney as a single nephrons split into three segments: the “proximal segment (ps)” which includes the proximal tubule and the loop of Henle, the “distal segment (dt)” that includes the distal convoluted tubule and the connecting tubule, and the collecting duct (cd).

Let *η*_pt−Kreab_ denote the fractional K^+^ reabsorption along the proximal tubule. In normal K^+^ intake this is about 67%. Fractional K^+^ reabsorption along the loop of Henle is denoted by *η*_LoH−Kreab_ and is about 25%. Therefore proximal segment reabsorption, denoted by *η*_ps−Kreab_, is given by

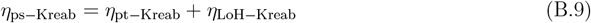

so that net proximal segment K^+^ reabsorption is given by

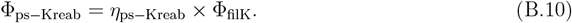

Distal tubule K^+^ secretion is modeled using a baseline values of 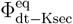 and is regulated by ALD and the gastrointestinal feedforward mechanism so that

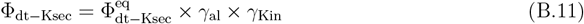

where *γ*_al_ denotes the regulatory effect of ALD (see Section B.3) and *γ*_Kin_ represents the gastroin-testinal feedforward effect (see Section B.5). Similarly, collectiong duct K^+^ secretion (Φ_cd−Ksec_ has a baseline value of 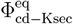 and is regulated by ALD so that:

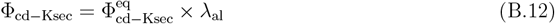

where *λ*_al_ denotes the regulatory effect of ALD (see Section B.3). Collecting duct K^+^ reabsorption (Φ_cd−Kreab_) is modeled as a linear function depending on the filtrate entering the collecting duct based transport from the previous segments:

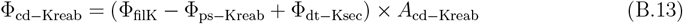

Urine K^+^ excretion (Φ_uK_) is given by the filtration and the net transport along the various segments:

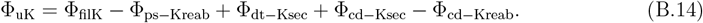

### B.3 Aldosterone effects

To model [ALD], denoted by *C*_al_, we use the approach developed by Maddah & Hallow [22]:

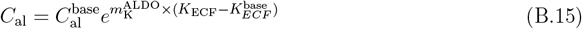

where 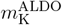 is a fitting parameter and *K*_ECF_ is the extracellular [K^+^] given by

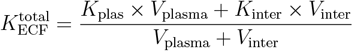

and 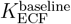 is the baseline extracellular [K^+^]. We capture the effect of [ALD] on Na^+^-K^+^-ATPase abundance by the scaling factor *ρ*_al_ (see Eq. B.6), which is represented linearly based on the findings of Phakdeekitcharoen et al. [30]:

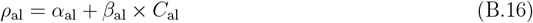

where *α*_al_ and *β*_al_ are parameters. The effect of [ALD] on distal tubule and collecting duct K^+^ secretion are represented by *γ*_al_ and *λ*_al_, respectively:

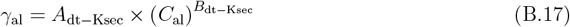

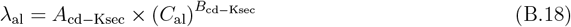

where *A*_dt−Ksec_, *B*_dt−Ksec_, *A*_cd−Ksec_ and *B*_cd−Ksec_ are parameters.

### B.4 Insulin effects

The concentration of plasma insulin, denoted by *C*_insulin_ in the time after a meal is given by

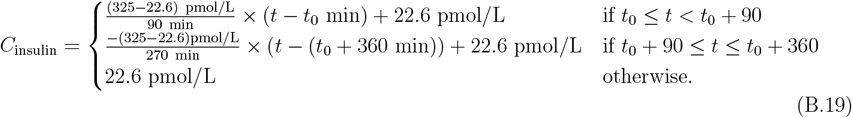

where *t*_0_ is the time at the beginning of the meal. To model insulin stimulation of Na^+^K^+^-ATPase, we let *ρ*_insulin_ (see (B.6)) be given by

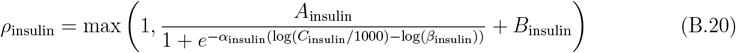

where *A*_insulin_ and *B*_insulin_ are fitting parameters.

### B.5 Gastrointestinal feedforward effect

The model represents gastrointestinal feedforward effect via the term *γ*_Kin_, which alters distal tubule K^+^ secretion depending on *M*_Kgut_ (see Eq. B.11) where

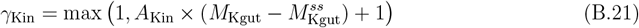

where *A*_Kin_ is a fitting parameter and 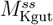 is the steady state value of *M*_Kgut._

### B.6 Parameters

The model parameter values and ranges used in the Morris analysis are given in Table B.1. Parameter fitting is discussed in Ref. [39].

**Table B.1:** Parameter ranges and baseline values (from Ref. [39]) used for the Morris analysis and model simulations. Parameters without a range given were not used in the Morris analysis.

| Name | Range | Original | Description |
| --- | --- | --- | --- |
| $V_{\text{plasma}}$ | — | 4.5 | plasma volume |
| $V_{\text{inter}}$ | — | 10 | interstitial volume |
| $V_{\text{intracellular}}$ | — | 24 | intracellular volume |
| $k_{\text{gut}}$ | 0.5e-2 - 1.5e-2 | 1.0e-2 | $\text{K}^+$ delivery rate to plasma |
| $P_{\text{trans}}$ | — | 0.78 | effective $\text{K}^+$ permeability from intracellular fluid |
| $K_{\text{m}}$ | — | 1.4 | half maximal activation of NKATPase |
| $V_{\text{max}}$ | — | 130 | maximum $\text{K}^+$ flow into cells |
| $P_{\text{ECF}}$ | — | 0.3 | effective $\text{K}^+$ permeability in extracellular fluid |
| $A_{\text{Kin}}$ | 0.13 - 0.38 | 0.25 | parameter that determined $\gamma_{\text{Kin}}$ |
| $\Phi_{\text{GFR}}^{\text{base}}$ | 0.1 - 0.13 | 0.12 | baseline glomerular filtration rate |
| $\eta_{\text{pt-Kreab}}^{\text{base}}$ | 0.34 - 0.74 | 0.67 | baseline fractional PT reabsorption |
| $\eta_{\text{pt-Kreab}}$ | 0.18 - 0.74 | 0.36 | fractional PT reabsorption (high $\text{K}^+$ diet) |
| $\alpha_{\text{TGF}}$ | 0.058 - 0.18 | 0.12 | TGF parameter |
| $\eta_{\text{LoH-Keab}}$ | 0.125 - 0.3 | 0.25 | loop of Henle fractional reabsorption |
| $\Phi_{\text{dt-Ksec}}^{\text{eq}}$ | 0.021 - 0.062 | 0.041 | baseline distal tubule $\text{K}^+$ secretion |
| $A_{\text{dt-Ksec}}$ | 0.17 - 0.52 | 0.35 | parameter that determines $\gamma_{\text{al}}$ |
| $B_{\text{dt-Ksec}}$ | 0.12 - 0.36 | 0.24 | parameter that determines $\gamma_{\text{al}}$ |
| $\Phi_{\text{cd-Ksec}}^{\text{eq}}$ | — | 2.2e-3 | baseline collecting duct $\text{K}^+$ secretion |
| $A_{\text{cd-Ksec}}$ | — | 0.16 | parameter that determines $\lambda_{\text{al}}$ |
| $B_{\text{cd-Ksec}}$ | — | 0.41 | parameter that determines $\lambda_{\text{al}}$ |
| $A_{\text{cd-Kreab}}$ | 0.25 - 0.75 | 0.50 | parameter that determines $\Phi_{\text{cd-Kreab}}$ |
| $A_{\text{insulin}}$ | 0.5 - 1.5 | 1.0 | parameter that determines $\rho_{\text{insulin}}$ |
| $B_{\text{insulin}}$ | 0.33 - 1.0 | 0.66 | parameter that determines $\rho_{\text{insulin}}$ |
| $K_{\text{IC}}^{\text{base}}$ | — | 130 | baseline intracellular concentration |
| $K_{\text{ECF}}^{\text{base}}$ | — | 4.2 | baseline extracellular concentration |
| $C_{\text{al}}^{\text{base}}$ | 42.5 - 127.5 | 85 | baseline ALD concentration |
| $m_{\text{K}}^{\text{ALDO}}$ | 0.25 - 0.75 | 0.5 | parameter that determines $C_{\text{al}}$ |
| $\alpha_{\text{al}}$ | 0.37 - 1.11 | 0.74 | parameter that determines $\rho_{\text{al}}$ |
| $\beta_{\text{al}}$ | 1.52e-3 - 4.6e-3 | 3.04e-3 | parameter that determined $\rho_{\text{al}}$ |

## C Sensitivity analysis

### C.1 Morris Method Description

Let *f* be a mathematical model where the output is given by *f* (*X*_1_, *X*_2_, …, *X*_*n*_) and *X*_1_, *X*_2_, …, *X*_*n*_ are the model inputs, which will be referred to as factors. When computing a sensitivity analysis using the Morris method, we start by determining a defined range for all the possible values for the factors. The factors are then rescaled to be uniformly distributed on the unit interval and an initial base value is selected at random from this distribution. Subsequently, one random factor, *X*_*i*_, is incremented by a step size Δ, typically chosen to be *n/*(2(*n* − 1)), where *n* is the number of factors. The elementary effect for the *i*-th factor, denoted by *EE*_*i*_ is computed by:

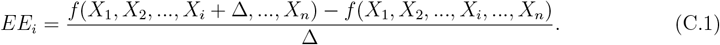

From this next value another random factor is incremented and the subsequent elementary effect is computed, until an elementary effect for each factor has been determined. This process is repeated *r* times by sampling at different points in the factor space. As a result, there will be a total of *r* elementary effects per factor at the end of the computations.

After computing the elementary effects for each factor, we can find the average value (*μ*_*i*_), average of the absolute values (*μ*^∗^), standard deviation (*σ*), and Morris Index (*MI*_*i*_):

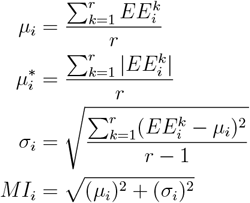

where 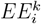 denotes the elementary effect of the *i*-th factor during the *k*-th model evaluation. The metrics can be interpreted in the following ways:

- *μ*^∗^ gives a factor ranking: a greater 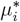 value indicates that the *i*-th factor affects the model output
- *σ* indicates that this factor interacts with other factors or is nonlinear
- *MI*: The Morris index is another way to factor rank by giving a single metric that incorporates both the mean (*μ*) and standard deviation (*σ*)

In an ordinary differential equation (ODE) model scenario, the factors are typically the model parameters. The model output measured can taken as the steady state solution, the values of the state variables at given time points in a simulation, or another measure that can be computed from a given model simulation. In this study, the model output was the final plasma and intracellular [K^+^] after a 50 simulation under high K^+^ intake. The ranges for the parameters used in the Morris analysis are given in Table B.1.

